# A systems and treatment perspective of models of influenza virus-induced host responses

**DOI:** 10.1101/368241

**Authors:** Ericka Mochan, Emily Ackerman, Jason E. Shoemaker

## Abstract

Severe influenza infections are often characterized as having unique host responses (e.g. early, severe hypercytokinemia). Neuraminidase inhibitors can be effective in controlling the severe symptoms of influenza but are often not administered until late in the infection. Several studies suggest that immune modulation may offer protection to high risk groups. Here, we review the current state of mathematical models of influenza-induced host responses. Selecting three models with conserved immune response components, we determine if the immune system components which most affect virus replication when perturbed are conserved across the models. We also test each model’s response to a pre-induction of interferon before the virus is administered. We find that each model emphasizes the importance of controlling the infected cell population to control viral replication. Moreover, our work shows that the structure of current models does not allow for significant responses to increased interferon concentrations. These results suggest that the current library of available published models of influenza infection does not adequately represent the complex interactions of the virus, interferon, and other aspects of the immune response. Specifically, the method used to model virus-resistant cells may need to be adapted in future work to more realistically represent the immune response to viral infection.

## 1. Introduction

Influenza A virus (IAV) leads to acute respiratory disease and significant morbidity and mortality around the world each year; the World Health Organization estimates 3 to 5 million cases of severe illness and 300,000-650,000 deaths worldwide every year are caused by IAV [1]. Generally, severe outcomes are limited to high-risk patient groups, i.e. infants, aged adults, or individuals with compromised immune systems. Occasionally, however, new strains emerge with pandemic potential that can induce severe disease across a broad portion of the population. For example, the 1918 Spanish influenza pandemic is estimated to have been responsible for the death of 2% of the world’s population between 1918 and 1920 [2]. Several pandemics have occurred since, including outbreaks in 1957, 1968, and 2009 [3,4]. Experts believe that avian H5N1 influenza viruses pose the greatest risk to public health. H5N1 infections have demonstrated the ability to cause severe disease in humans, including symptoms such as fever, respiratory symptoms, lymphopenia, and cytokine storm (hypercytokinemia) [5–7]. Cytokine storm occurs when the host experiences out-of-control pro-inflammatory responses and insufficient anti-inflammatory responses to infection. This is often a result of severe influenza infection and causes acute respiratory distress syndrome (ARDS) and multiple organ failure in many patients [7].

Often, IAV infections are treated with neuraminidase inhibitors, such as oseltamivir (i.e. TamiFlu), which can be highly effective if administered during the early infection phase. However, IAV-infected hosts often do not seek treatment until late in their infection when the virus is already present at high levels and it may be too late for an effective treatment. Especially in the case of H5N1, neuraminidase inhibitors are often ineffective at containing cytokine storm and do not prevent the excess morbidity and mortality seen in these infections [8,9]. Moreover, oseltamivir-resistant strains can quickly evolve, as observed during the 2009 H1N1 pandemic [8].

### 1.1 Immune modulation for the treatment of IAV infection

Modulating the immune response post infection to control inflammation or pre-infection to provide increased protection for high risk groups has been a major theme in severe influenza infection research [10–20]. Corticosteroids have been suggested as a potential treatment option for patients undergoing severe IAV infection with accompanied cytokine storm, while pre-stimulating interferon-associated pathways have been suggested to protect high-risk groups [7,20–22]. Corticosteroids have anti-inflammatory effects on the host and thus may be used to treat hypercytokinemia. The impact of corticosteroids on IAV-infected hosts is currently inconclusive. While some studies indicate steroids are effective in alleviating influenza symptoms in human patients [23,24], others have shown increased mortality in patients treated with corticosteroids [25–27].

Interferon (IFN) is a key regulator of the innate immune system, and pre-stimulation of interferon regulating pathways has provided a preventative advantage in mice infected with deadly influenza viruses [21,22,28]. IFN is essential for viral clearance, has been heavily studied since its discovery in 1957 [29], and has a complicated role in immunopathology (See [30]for a review). In recent mouse studies, animals, prior to infection, were exposed to synthetic or natural agonists of the Toll-Like Receptor pathways (specifically TLR3 and TLR4) that activate IFN production [22,28,31,32]. This pre-stimulation induced higher concentrations of IFN in lung epithelial cells, reduced virus titers and significantly improved infection outcomes in animals infected with highly pathogenic viruses. Interestingly, some studies have shown that select bacterial strains in yogurt provide protection against influenza infection by increasing IFN production [33,34]. The suggested mechanism is that exopolysaccharides produced by the bacteria exert immunostimulatory effects via the TLR pathways. These evidences combined with the several studies demonstrating dysregulation of the immune response during deadly influenza infections [13,17,35,36] suggests that immunomodulation prior to infection may be an option for protecting high risk groups. Moreover, as IFN is a common component of mathematical models of influenza-induced immune responses, the ability to replicate the effects of pre-stimulating IFN-regulating pathways provides a valuable measure of model applicability.

### 1.2 Mathematical models of the lung host response to IAV infection

Mathematical models of the immune response in IAV-infected lungs have previously been used as a computational platform for treatment optimization [37–43]. Modeling can be an invaluable tool for ascertaining kinetic parameters of an influenza infection which are difficult to measure in traditional experiments. Many experimental data sources, particularly murine (mouse) models of influenza, are generated from a pool of measures collected from hosts subjected to identical experimental conditions. Multiple hosts are sacrificed at pre-determined intervals and measured for variables of interest. Because these animals need to be sacrificed to measure cell and cytokine levels, hosts cannot be tracked for the full duration of the infection, making true longitudinal data impossible to obtain. These experiments assume that the all animals will react nearly identically to the infection, but inter-individual variability in hosts can invalidate this assumption. Mathematical modeling can be used to help fill in gaps in knowledge created by the deficiencies in experimental data. Models can vary substantially in complexity, depending on the facets of the immune response they contain and the number of interactions represented.

Models generally fall into one of two categories: target cell-limited models, in which the healthy epithelial cells, which act as a target for the virus, are unable to replicate themselves [38,40], or models in which healthy cells are able to regenerate [37,39,41–43]. These models all feature three basic components: healthy epithelial cells, infected epithelial cells, and the virus. More components, such as cytokines, immune cells, or antibodies, can be added to the model with additional equations and parameters. Larger models can provide a more comprehensive understanding of the immune response but may also require a larger pool of data from which to calibrate the model. Smaller models do not need as much data for training the model, but they may also make more simplifying assumptions that can be difficult to support biologically.

In this work, we review three recently published models that contain similar components of the immune response. We consider two points of comparison. The “systems” perspective being which components of the model most strongly regulate virus replication. And the “treatment” perspective being how well do the models recapitulate the observations in IAV-infected animals whose immune system has been stimulated prior to infection. Pre-stimulation is simulated by increased concentration of IFN prior to infection. We find that virus concentration is largely influenced by the proportion of cells in the model that can become infected or virus-resistant; therefore, controlling these populations is paramount to controlling viral replication. We also find that current models do not capture the effect of increased IFN concentrations on suppressing virus replication.

## 2. Description of IAV Immune Response Models

To provide a review of current models of IAV-induced immune responses, we selected recent models which contain common elements of the innate immune response. This allowed for easier comparisons between model analyses. The models analyzed are: Saenz et al. 2010 [38], Pawelek et al. 2012 [37], and Hancioglu et al. 2007 [39]. Figure 1 depicts the interactions represented within each of the models. The Saenz and Pawelek models are trained to experimental data (e.g. cytokine concentrations and immune cell counts) measured in pony lungs infected with H3N8 virus, while the Hancioglu model was fit to certain qualitative behaviors selected from a study of the human response to IAV infection by Bocharov and Romanyukha [44].

**Figure 1.**
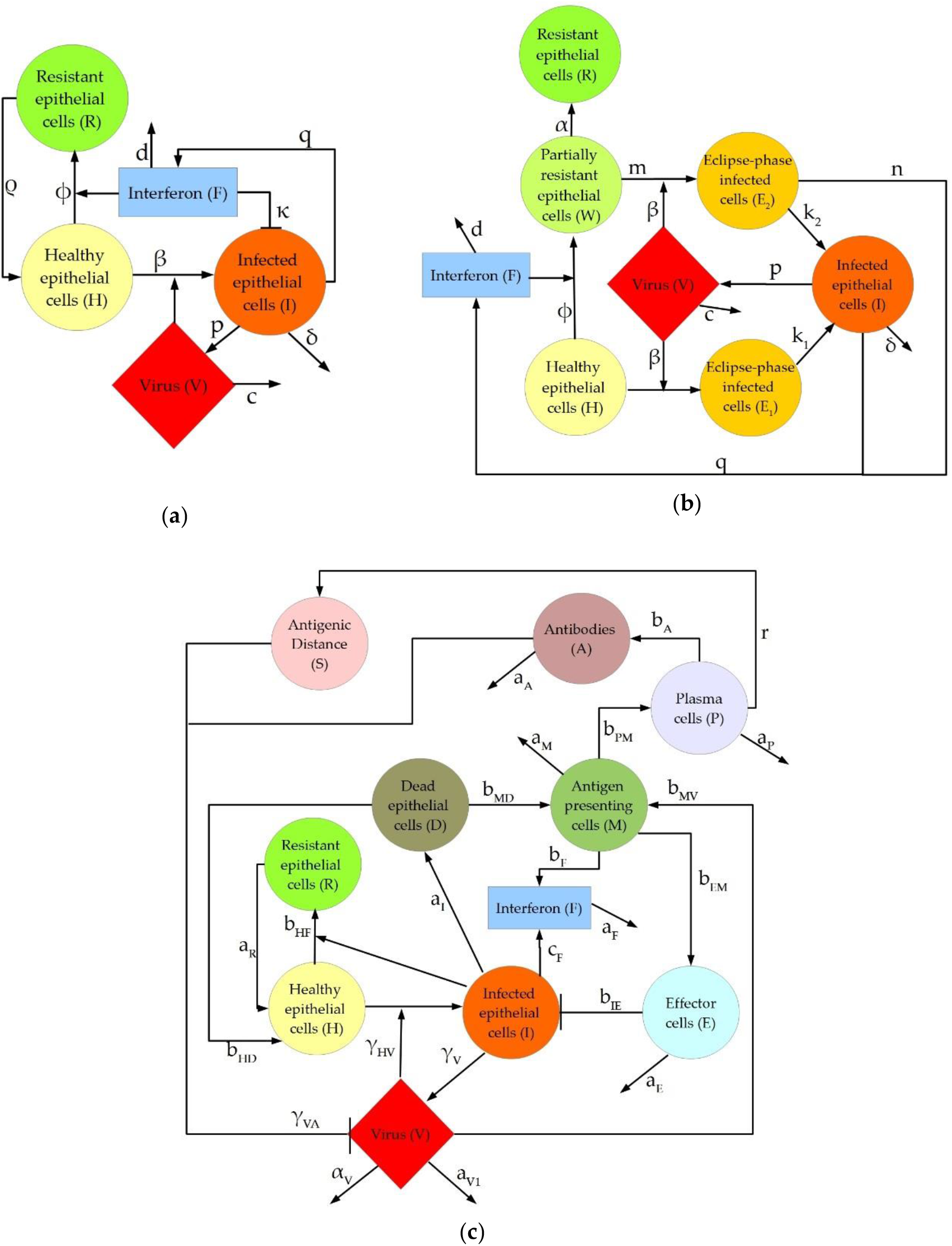
Graphical depiction of interactions represented in the three models discussed in this manuscript: (**a**) Pawelek model; (**b**) Saenz model; (**c**) Hancioglu model.

Five elements of the intrahost immune response are conserved across each model: healthy epithelial cells (H), infected cells (I), virus (V), type I interferon (F), and “resistant cells”, i.e. epithelial cells with interferon-induced virus resistance (R). While each model has these five features in common, the inflammatory response to viral infection is represented differently, depending largely on model complexity. These differences are particularly apparent in the model-specific incorporation of the production, activity, and depletion of IFN. In the Pawelek model (Figure 1a), interferon has two functions: creating virus-resistant cells when interacting with healthy epithelial cells, and increasing infected cell death when interacting with infected epithelial cells. In the Saenz model (Figure 1b), interferon leads to the creation of virus-resistant cells but does not impact the infected cells directly. Instead, the infected cells produce more interferon. The Hancioglu model (Figure 1c) uses interferon to create resistant cells (as in the other two models) while interferon is produced by infected cells and antigen-presenting cells. In all models, a decrease in interferon levels is caused by a combination of natural decay and absorption into epithelial cells.

## 3. Materials and Methods

Three ordinary differential equation (ODE) models of the intrahost immune response to IAV infection that explicitly included type I interferon were chosen from literature. These three published models were selected for their significant variance in complexity; specifically, in the interactions of IFN with other model components. For each model, the immune response is simulated in MATLAB version R2017a using the parameter values and initial conditions published in the original papers. Integration was performed with ode23s.

We performed two main assessments on the three featured models: a local sensitivity analysis and an interferon pre-stimulation study. Sensitivity analysis was performed using a MATLAB package previously published by Nagaraja et al [45]. The Param_var_local.m function performed a local sensitivity analysis on the virus equation in each model to all parameters over a ten-day simulation. The function increases and decreases each parameter in the model by 1% and recalculates the solution to the system of ODEs. Sensitivity is then calculated with the central finite difference formula to generate logarithmic sensitivities of each equation to each parameter in the model. The sensitivity of each parameter was ranked by the area under the curve (AUC). Parameters which yield the highest AUC over the full ten-day simulation are judged to be the most sensitive.

Two tests were used to evaluate each model’s reaction to simulated interferon pre-stimulation. First, four values of the initial level of the IFN present in the system (F0) were tested to assess whether increased initial IFN levels will inhibit viral growth, peak, or clearance. In each case, while the initial condition on the IFN equation changed, all other initial conditions and parameters remain constant. Additionally, the amount of time between the initial IFN induction and the start of the infection was varied by delaying the onset of the virus infection with respect to the IFN. In all cases, induction of IFN via IFN-regulating pathways is modeled as a step change in IFN concentration. The 6 possible delays in the virus administration included 0, 2, 4, 6, 8, and 10 days (equivalent to pre-stimulating IFN 0, 2, 4, 6, 8, and 10 days prior to infection). The initial level of IFN is kept at 1 in these simulations, though the published initial condition of the fold change of IFN in the Saenz model is 0. To simulate a true pre-stimulation, there must be a nonzero initial level of IFN to observe the impact of IFN on the remainder of the system.

## 4. Results

### 4.1. Sensitivity analysis

#### 4.1.1. Pawelek et al. model

The Pawelek model includes five equations and nine parameters to model the intrahost immune response to IAV infection. Epithelial cells begin as healthy target cells (H) and can become infected (I) through direct interaction with the virus (V) or can become resistant to infection (R) through interaction with type I interferon (F). Interferon can also lead to enhanced infected cell clearance by stimulating the activation of natural killer cells, which are not explicitly represented in the model. The structure of the Pawelek model is given below in equations 3.1.1, and a full list of parameters, their biological interpretations, nominal values, and units is provided in the Appendix in Table A1. Note that we used the original version of the equations presented in the paper and not the alternative version presented by Pawelek *et al*. which incorporates a time-varying death rate δ(t) for infected cells in only the later portion of the simulation. We did not use this artificial delay in infected cell clearance, as a delay equation would not be consistent with the other two models presented in this review. Instead, δ = 2/day for the entirety of the simulation.

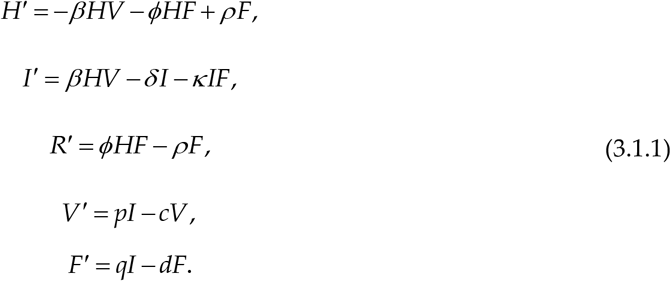

First, a local sensitivity analysis was used to identify model parameters to which the virus titer is most sensitive. The parameters which most affect the virus level change over time, as shown in Figure 2a. Over the first two days, the decay rate of interferon (d) and the production of resistant cells (φ) dominate the behavior of the virus. In the later phase of infection, virus behavior is controlled predominantly by the rate at which cells lose their resistance (ϱ), the death of infected cells (δ), and the depletion of interferon (d). This implies that in all stages of infection, virus levels are largely impacted by interferon and resistant cell populations. Given that interferon controls the number of resistant cells in the system, and thus the number of cells available to become infected by the virus, the production and depletion of interferon is expected to be vital to the control of the virus growth throughout the simulation. Over ten days, the parameters with the highest total AUC are the rate of loss of resistance in the epithelial cells (ϱ) and the decay rate of infected cells (δ). Table 1 shows the AUC for each parameter after a ten-day simulation.

**Figure 2.**
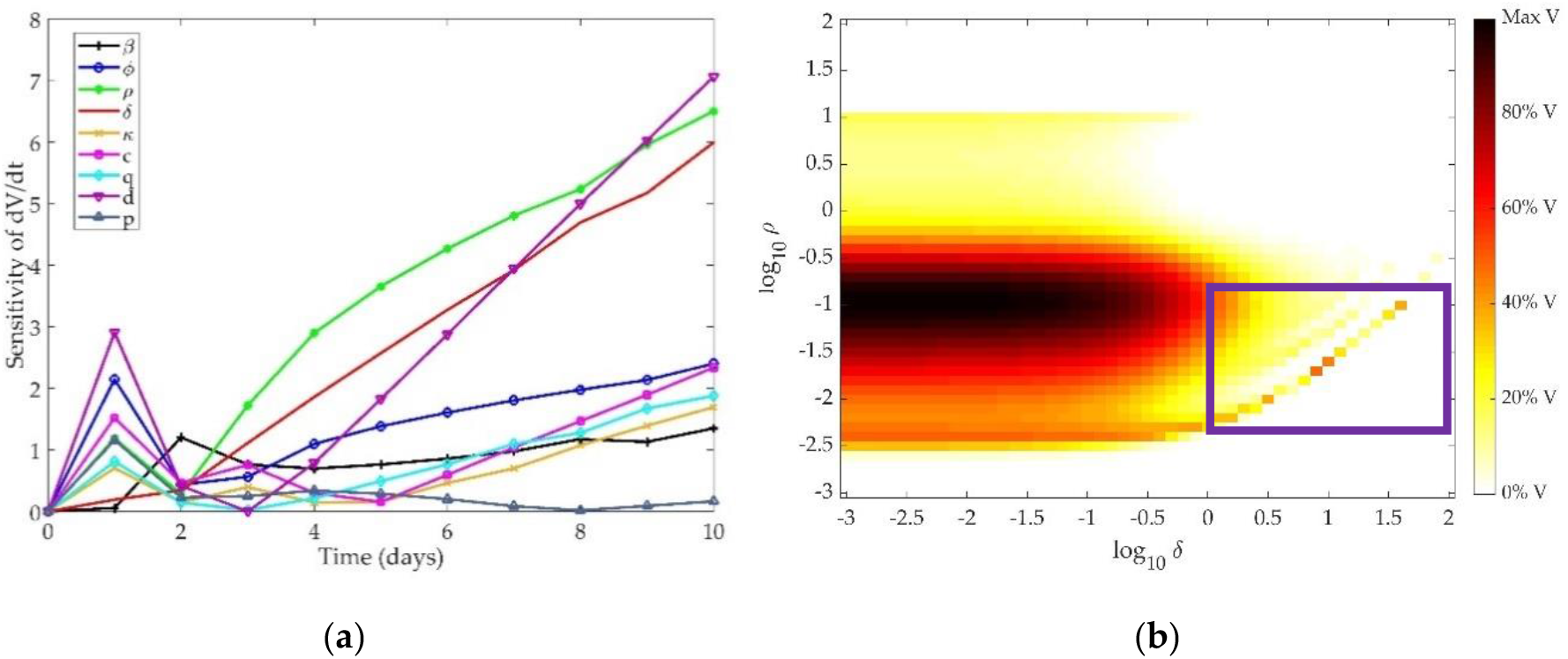
(**a**) Time-dependent sensitivity of virus to each parameter in the Pawelek model; (**b**) Two-dimensional sensitivity of Pawelek model to two most sensitive parameters: ρ and δ. Colors correspond to the amount of virus present at ten days post-infection in simulation as a fraction of the maximum virus concentration observed. Darker colors correspond to a higher level of virus present at day 10, while lighter colors indicate a low level of virus present in the system after ten days.

**Table 1.** AUC of virus equation for each parameter in the Pawelek model.

| Parameter | AUC |
| --- | --- |
| $\varrho$ | 36.87 |
| $\delta$ | 29.43 |
| $d$ | 24.83 |
| $\phi$ | 14.93 |
| $\beta$ | 8.68 |
| $q$ | 6.59 |
| $c$ | 4.37 |
| $\kappa$ | 4.01 |
| $p$ | 1.90 |

Figure 2b depicts the model response to changing these two most sensitive parameters concurrently. Colors on the graph correspond to the amount of virus present in the system after ten days, where darker colors indicate more virus present. As δ increases, the infected cells die at a faster rate, and the virus cannot be sustained, leading to a lower virus level at the end of the ten-day simulation. As ρ increases, the healthy epithelial cells regenerate at a faster rate, providing a larger pool of cells which may transition to infected cells and produce virus. Thus, as ρ increases, more virus is expected to be present at the end of the simulation. However, the current formulation of the model may overfit the parameters, leading to unexpected trends in the end behavior of the virus. Specifically, similar values of ρ and δ result in highly different responses (see lower right portion of Figure 2b, where log(δ) = 0 to 2 and log(k_2_) = −2.5 to −1, demonstrated in the purple box). As more cells become resistant, there is greater feedback to the healthy cell population, allowing for a slight rebound in the target cell population. With more target cells available to become infected, there is an increase in virus over time. The structure of the model thus results in the creation of virus over time with the addition of more cells that are resistant to infection. This is an oversimplification of the interaction of interferon with epithelial cells and is likely not a true representation of intrahost dynamics.

#### 4.1.2. Saenz et al. model

The Saenz model includes eight equations and twelve parameters. The epithelial cell population is divided into six subtypes, based on whether these cells have been infected with virus (V) and/or affected by type I interferon (F). Cells can be healthy cells (H), infected cells that are unprotected by interferon (E1), partially resistant healthy cells (W), infected cells that have been affected by interferon (E2), productively infected cells (I), or fully resistant cells (R). A list of the parameters, their biological interpretations, nominal values, and units is provided in the Appendix in Table A2. The structure of the Saenz model is given below in equations 3.1.2:

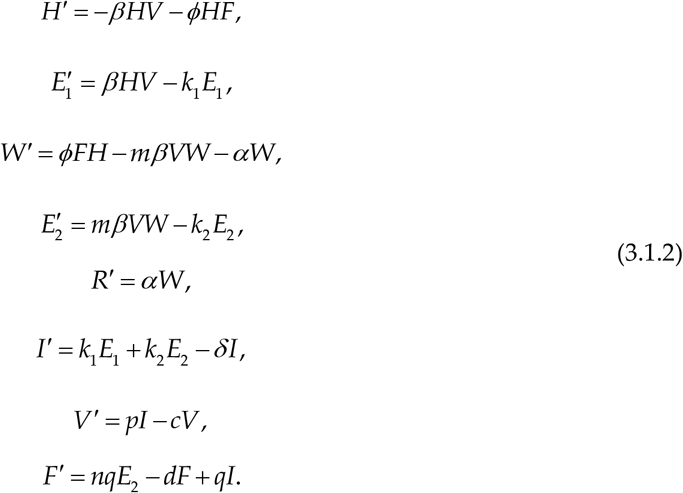

We explored the time-dependent sensitivity of the virus to each parameter in the model (Figure 3a). The first three days post-infection are predominantly controlled by the infectivity of the virus (β), production of virus by infected cells (p), and creation of unprotected infected cells (k_1_). The later phase of the infection is controlled by death of infected cells (δ) and the eclipse phase period of interferon-protected infected cells (k_2_). Table 2 shows the AUC for each parameter after a ten-day simulation.

**Figure 3.**
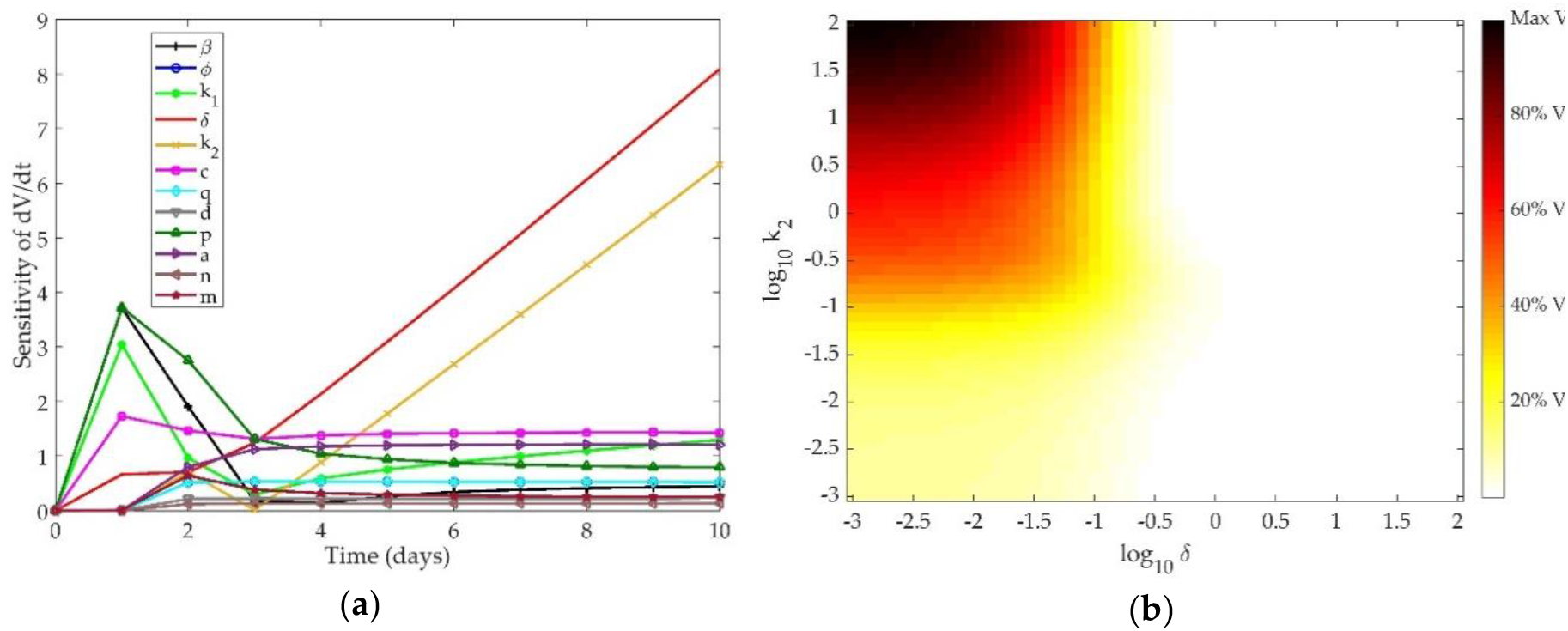
(**a**) Time-dependent sensitivity of virus to each parameter in the Saenz model; (**b**) Two-dimensional sensitivity of Saenz model to two most sensitive parameters: k_2_ and δ. Colors correspond to the amount of virus present at ten days post-infection in simulation as a fraction of the maximum virus concentration observed. Darker colors correspond to a higher level of virus present at day 10, while lighter colors indicate a low level of virus present in the system after ten days.

**Table 2.** AUC of virus equation for each parameter in Saenz model.

| Parameter | AUC |
| --- | --- |
| $\delta$ | 38.26 |
| $k_2$ | 24.50 |
| $c$ | 14.43 |
| $p$ | 13.86 |
| $a$ | 10.40 |
| $\phi$ | 4.75 |
| $q$ | 4.75 |
| $\beta$ | 3.35 |
| $k_1$ | 3.07 |
| $m$ | 2.93 |
| $d$ | 2.00 |
| $n$ | 1.18 |

As in the Pawelek model, controlling the death rate of infected cells indirectly controls the death rate of the virus, as infected cells are the only source of free virus in the system. The eclipse phase also indirectly controls the virus production, since infected cells begin producing free virus as soon as the eclipse phase ends and become productively infected cells. The parameters most closely related to IFN concentration in the model (n, q, d, and φ) are not among the most sensitive for the virus in this model.

A two-dimensional scan of the most sensitive parameters to the clearance rate of the virus from the host is shown in Figure 3b. As expected, when k_2_ and δ are high, the infected cells die at an increased rate and the virus is completely cleared from the host after ten days. When these parameters are low, however, the infected cell population is sustained and the virus remains at high levels ten days post-infection.

#### 4.1.3. Hancioglu et al. model

The Hancioglu model is the largest and most complex of the three models, featuring ten equations and 29 parameters. The model includes healthy (H), resistant (R), and infected (I) epithelial cells, virus (V), and interferon (F), as in the previous models. In addition, macrophages (M), T cells (E), plasma cells (P), antibodies (A), and antigenic distance (S) are incorporated, as shown in Figure 1c. The total number of epithelial cells is assumed to be constant, so the number of dead epithelial cells (D) does not require a differential equation, but rather obeys an algebraic formula: *D* = 1 − *H* − *I* − *R*. Parameters, biological definitions, and their nominal values are defined in Table A3 in the Appendix. Note that all parameters in this model are scaled such that they are unitless. The equations which define the Hancioglu model are given below in (3.1.3):

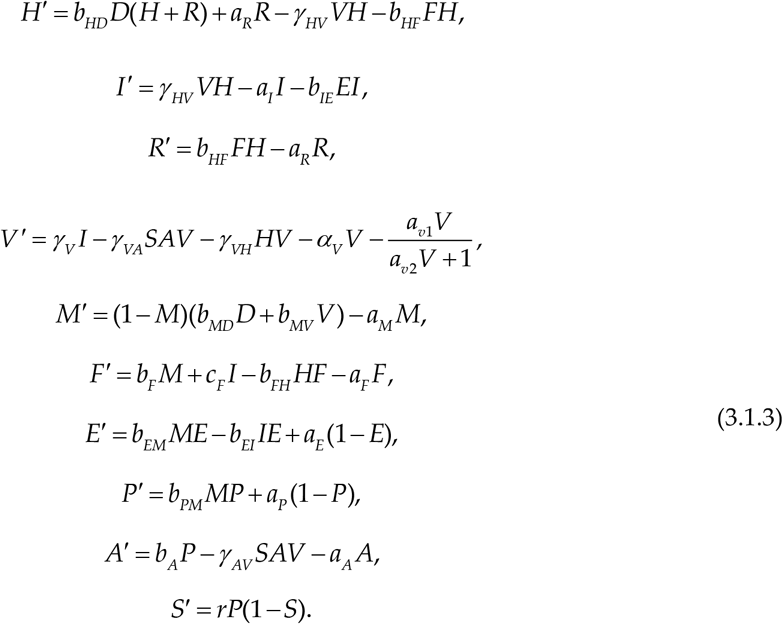

Figure 4a shows the sensitivity of the virus equation to each of the parameters of the model. Table 3 gives the total AUC for each parameter in the Hancioglu model over the ten-day simulation.

**Figure 4.**
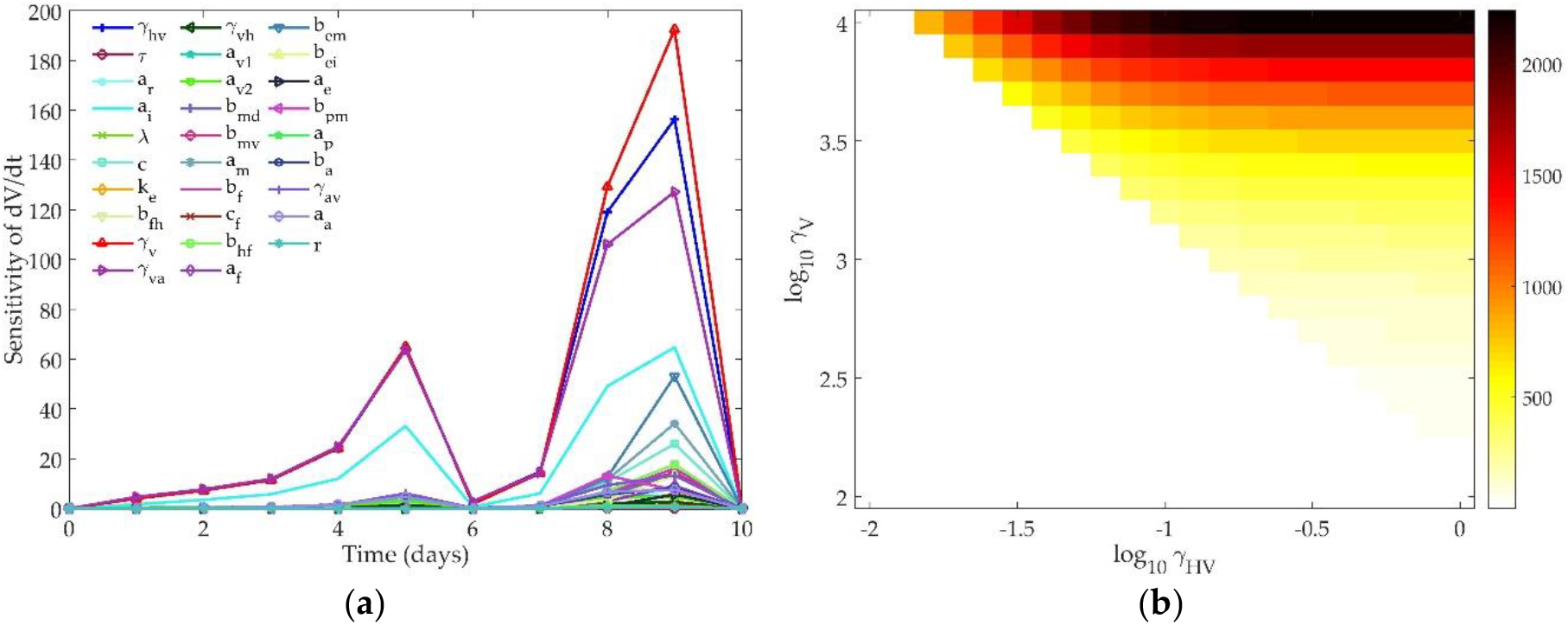
(**a**) Time-dependent sensitivity of virus to each parameter in the Hancioglu model; (**b**) Two-dimensional sensitivity of Hancioglu model to two most sensitive parameters: γvh and γv. Colors correspond to the maximum amount of virus present over a ten-day simulation. White indicates that the maximum value of V is 0.01, the initial value of the virus. Darker colors indicate higher values of peak virus.

**Table 3.**
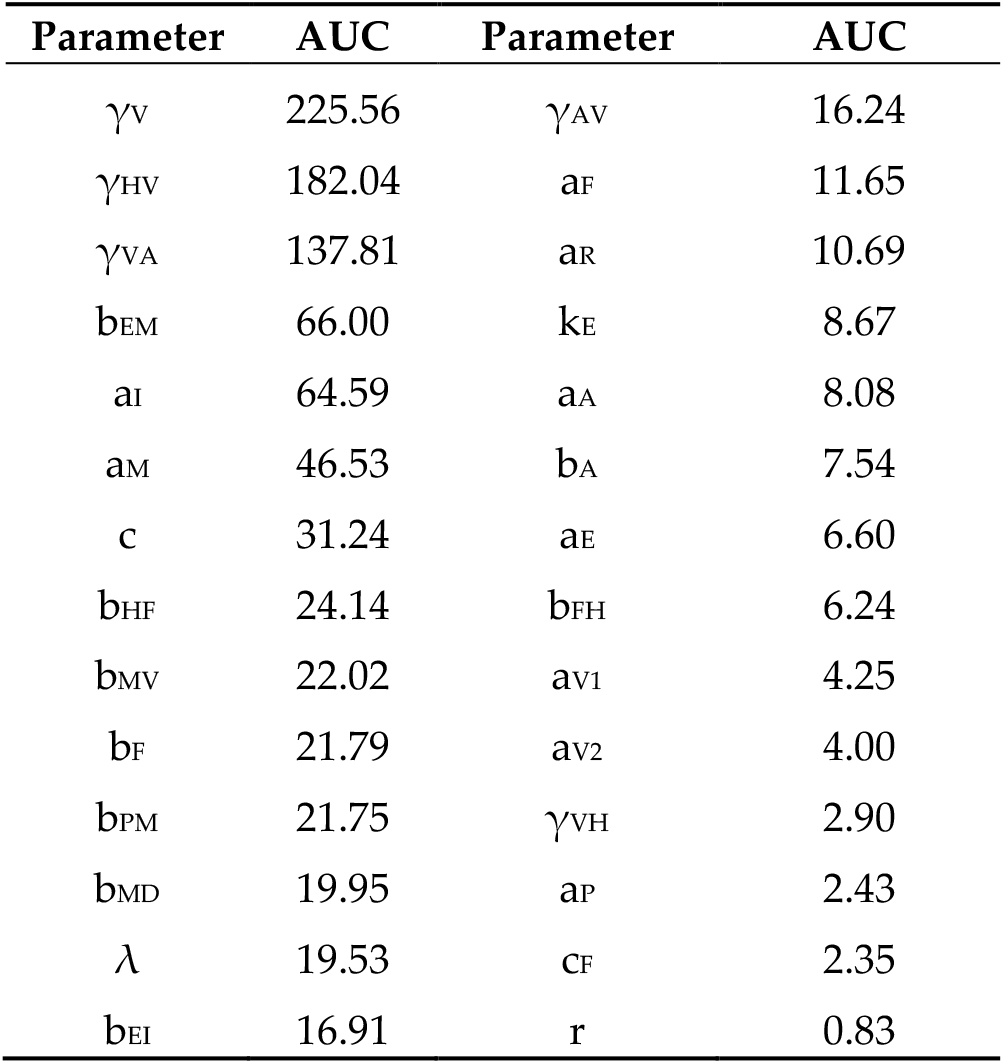
AUC of virus equation for each parameter in Hancioglu model.

The early stage infection is largely controlled by virus production by infected cells (γv), infected cell death (ai), and loss of resistance in epithelial cells (ar). All three of these parameters are essential to controlling the infected cell population, which, like in the previous two models, controls the amount of free virus produced by the host. In the later stages of infection, γv and the infectivity of the virus (γhv) are most influential on the virus trajectory.

The amount of virus present at the end of the simulation does not change based on γv or γhv; the model structure will lead to complete clearance of the virus even if these parameters have changed by several orders of magnitude. The mechanism by which the virus will clear does change based on the values of γv and γhv, as the peak value of the virus is impacted by these parameters. Figure 4b shows the change in the peak value of the virus as these two most sensitive parameters are changed concurrently. The nominal model creates a peak virus level of about 100. Varying these parameters can lead to a much higher peak viral titer. Changing γv or γhv can create a bifurcation in the virus behavior where the virus will either rise to its peak before being cleared by the immune response, or the virus will monotonically decay and will not instigate an immune response. When both γv and γhv are low (bottom left corner of Figure 4b), the infectivity and replication rate of the virus are too small to sustain the viral titers, and the virus will harmlessly decay to zero within a few days post-infection. As these two parameters increase, the virus replication is strong enough to sustain an infection and the immune response will be activated. Higher γv values cause the virus to peak faster, which then can lead to faster clearance, as the immune components in this model are virus-dependent (see Figure 1c).

### 4.2. Models’ response to simulated IFN pre-stimulation

We simulated IFN pre-stimulation in these models by changing the initial concentration of IFN in the system. Currently, several compounds which stimulate TLR pathways to induce IFN production and increase protection during severe respiratory infection are being studied [46]. The effect of these compounds on the dynamic response of the lung immune systems has not been determined. Several experiments have indicated a protective effect of pre-stimulation of TLR pathways before influenza infection in murine models [46–49]. We tested the three ODE models to determine whether they could replicate these experimental results. For each model, the response to changing both the magnitude of the initial interferon levels and the time between the initial interferon induction and the viral infection is reported.

#### 4.2.1 Pawelek model response to IFN pre-stimulation

Figure 5a depicts the impact of altering the initial levels of IFN (F0) present in the system when the virus is administered. The Pawelek model predicts three phases of behavior based on the amount of interferon concentration initially in the host. At high levels of interferon, there is a quick rise in the virus followed by a slow decay, leading to eventual clearance from the host. When F0 is 1, the virus exhibits biphasic behavior, with an initial peak approximately 12 hours post-infection and a second peak around 2 days post-infection, followed by slow clearance of the virus over the next week. When F0 is very low (less than 1), the same initial peak after 12 hours is followed by a long, slow rise in the virus trajectory after day 3. At high levels of F0, the virus peak is about 2 orders of magnitude lower than the other simulations, and the healthy cells show a slight rebound before dying out around 1 day post-infection. The IFN also monotonically decays when it is initially set to a high level and does not show the same rebound behavior seen in lower initial levels of IFN. Despite these three differences, it is difficult to determine how these rebounding virus trajectories would impact the survival of the host. The Pawelek model leads to complete death of all healthy cells in all trajectories, meaning there are no target cells remaining for the virus to infect after a few days of the infection. Because of this, the Pawelek model cannot accurately predict whether high initial concentrations of IFN would save the host.

**Figure 5.**
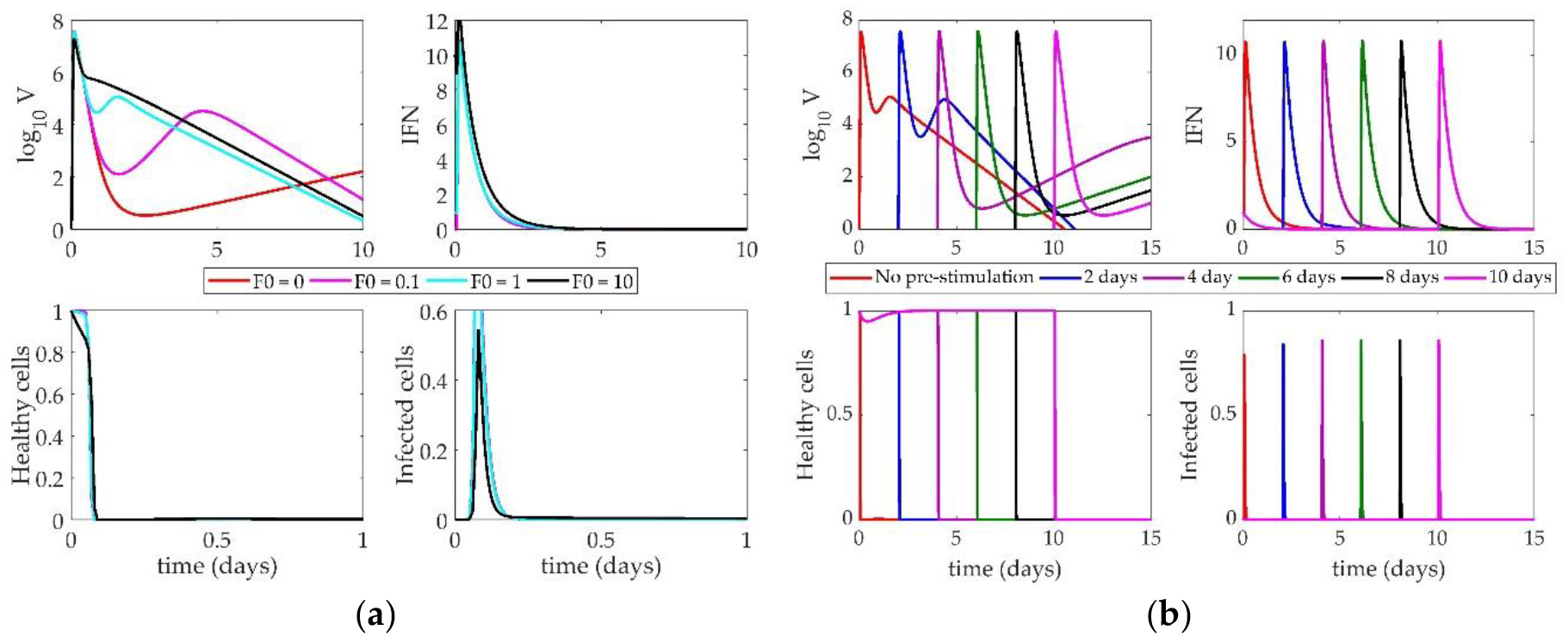
(**a**) Response of Pawelek model to increasing IFN on day 0. Lines correspond to different values of F0 (initial condition of interferon); (**b**) Impact of IFN pre-stimulation scheduling on the Pawelek model. IFN treatment is given on day 0 and patients are simulated to become infected 2, 4, 6, 8, or 10 days after treatment.

The impact of changing the time between the IFN pre-stimulation and the onset of the infection is shown in Figure 5b. Each line denotes a different simulation in which the virus is administered at the indicated day on the figure legend. Again, changing the time between treatment and infection causes three phases of behavior in the virus. When IFN induction and virus are given simultaneously (red “No pre-stimulation” line), the simulation describes the behavior of an untreated patient. In this case, the virus peaks almost immediately, falls, and then exhibits a smaller, secondary peak around day 2 as the infected cells have time to produce more virus.

The other simulations in Figure 5b show that the Pawelek model predicts a negative impact on the host after IFN pre-stimulation. With a 2-day pre-stimulation (blue line), the virus trajectory is approximately the same as the nominal model. The initial peak reaches the same magnitude as the nominal model, but the secondary peak is more pronounced. When IFN is given before the virus, there are no infected cells present in the system (which are the only sources of IFN in this model). Thus, the IFN levels will decrease until the virus is introduced on day 2. At that point, the IFN has decreased approximately two orders of magnitude, meaning the initial treatment was not sustained and has had a deleterious effect on the host, as there is less IFN present in the system than under normal circumstances.

As the delay between IFN pre-stimulation and virus lengthens, this effect becomes more pronounced. While the initial virus peak always reaches the same magnitude, the secondary peak, which controls the long-term behavior of the system, gets larger with the increased delay. This indicates that the model predicts a long-lasting influenza infection after significant IFN pre-stimulation rather than any improvement in patient outcomes, regardless of the length of time between virus and IFN treatment.

#### 4.2.2 Saenz model response to IFN pre-stimulation

The response of the Saenz model to an increase in the initial IFN present in the system is shown in Figure 6a. Changing the level of IFN creates a bifurcation in the virus peak magnitude. When F0 is below 1, the virus peak reaches 10^5^ PFU (a 5-fold change in the virus level). IFN and infected cells rise to their peaks around 3 days post-infection, and healthy cells die out within 2 days. When F0 is above 1, however, the virus peak only reaches approximately 10 PFU. IFN decays monotonically, and almost no infected cells are created. Healthy cells die out almost instantly upon infection, as more resistant cells can be created from the increased initial IFN levels.

**Figure 6.**
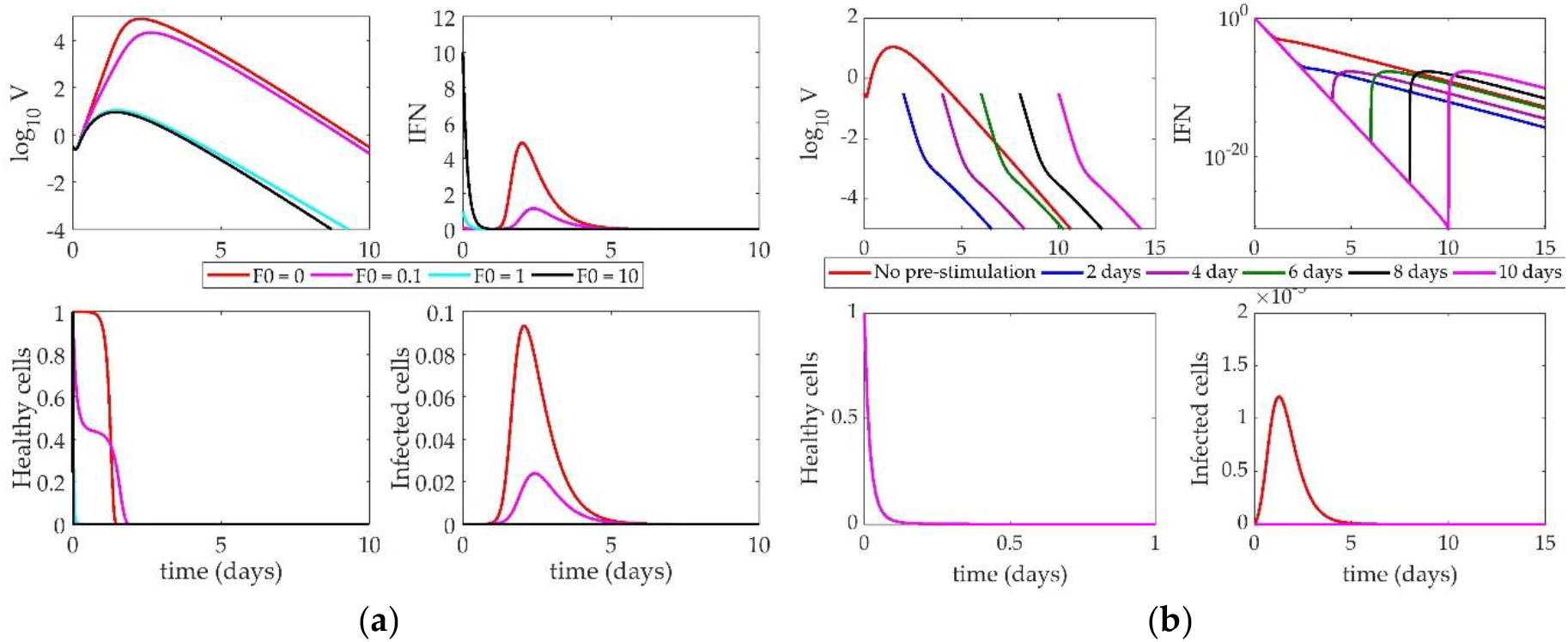
(**a**) Response of Saenz model to increasing IFN on day 0. Lines correspond to different values of F0 (initial condition of interferon); (**b**) Impact of IFN pre-stimulation scheduling on the Saenz model. IFN treatment is given on day 0 and patients are simulated to become infected 2, 4, 6, 8, or 10 days after treatment.

Figure 6b illustrates the impact of IFN pre-stimulation on the system. The nominal model (red “No pre-stimulation” line) is equivalent to the turquoise line (F0=1) in Figure 6a. As the time between the initial IFN pre-stimulation and virus infection increases, we see the pre-stimulation confers protection on the host by creating a larger pool of resistant cells initially, leading to a depletion of healthy cells within hours of the initiation of the simulation. When the host becomes infected, there is not a large enough pool of healthy cells available to create any substantial infected cell population. Without these cells, no new virus can be produced, so the virus monotonically decays to zero within only a few days post-infection.

Like the Pawelek model, the Saenz model cannot support high levels of IFN without the presence of virus, as only infected cells can produce more IFN. In all simulations, the IFN decays steadily and more rapidly than in the Pawelek simulations in Figure 5. This is largely due to the differences in the decay rate of IFN between the two models (Pawelek model, d = 1.9 day^−1^, Saenz model, d = 6.8 day^−1^).

#### 4.2.3 Hancioglu model response to IFN pre-stimulation

The Hancioglu model (Figure 7) shows very little sensitivity to the time of IFN induction or the magnitude of interferon concentration. There is a negligible difference in the time to the peak of the virus when F0 = 1 versus F0 = 100. The model’s parameters were fit in a way such that the behavior of the model does not change, even with an enormous initial influx of interferon. An initial absence of interferon has no impact on the virus trajectory. Despite the presence of resistant cells in the model, interferon has no major effect on the system.

**Figure 7.**
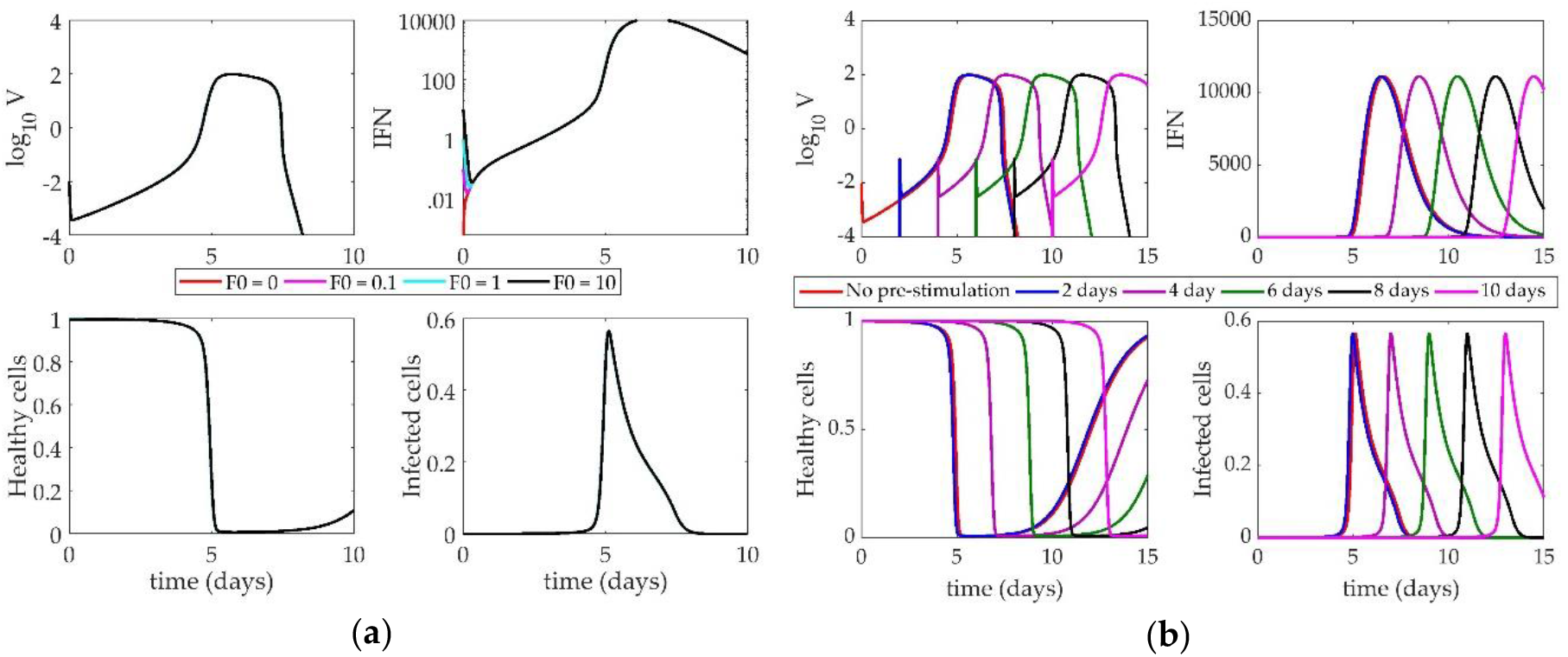
(**a**) Response of Hancioglu model to increasing IFN on day 0. Lines correspond to different values of F0 (initial condition of interferon); (**b**) Impact of IFN pre-stimulation scheduling on the Hancioglu model. IFN treatment is given on day 0 and patients are simulated to become infected 2, 4, 6, 8, or 10 days after treatment.

When the system is tested with a pre-stimulation of interferon (Figure 7b), there is little change in the behavior of the virus. The entire system shifts horizontally with the time delay, but the overall behavior of the model does not change. Like in previous models, without a starting virus population, the system cannot sustain the initial concentration of IFN. When the virus is finally introduced, the IFN level has essentially fallen to zero, making any impact from the pre-stimulation negligible.

## 5. Discussion

Interferon is known to have several antiviral effects in an IAV-infected host, including activating an antiviral state in epithelial cells, sensitizing cells to apoptosis, activating NK cells, and initiating the differentiation of cytotoxic T cells [50–53]. Each analyzed model represents a distinct subset of these interactions, including the creation and depletion of virus-resistant cells. In the Pawelek model, IFN is only produced by infected cells, and healthy cells can become resistant through an interaction with IFN. This resistance fades over time and cells return to a susceptible state. Resistant cells in the Hancioglu model also become susceptible, but IFN can be produced by either infected cells or antigen presenting cells. Conversely, the Saenz model features epithelial cells that are either partially or fully resistant to infection, and cells do not lose resistance over time.

Each of the three models shows a sensitivity of the virus to the creation and loss of infected epithelial cells. The virus equation of the Pawelek model is most sensitive to the loss of resistance in epithelial cells and the death rate of infected cells. If the infected cells die off too quickly, the virus cannot replicate at a rate high enough to sustain the infection. Similarly, if cells are becoming virus-resistant too quickly, there will not be a sufficient number of cells remaining to become infected and keep the viral titers elevated. In this way, the presence of the virus in the system is predominantly driven by the number of cells currently infected or able to become infected. The Saenz model also emphasizes a low death rate of infected cells, as well as a short eclipse phase for infected cells. The duration of the eclipse phase determines the delay in time between the infection of the cell and the subsequent release of virion by the infected cell. The shorter the eclipse phase, the more readily the cells can begin producing virus. As in the Pawelek model, the Saenz model shows that the availability of productively infected cells is vital to the continuation of the infection.

The Hancioglu model also emphasizes the importance of maintaining a large pool of infected cells, but through a different set of parameters than the Pawelek or Saenz models. The infectivity of the virus and the replication rate of the virus are the most sensitive parameters in the model. The Hancioglu model is thus controlling the virus by a high rate of production of infected cells, and not through a diminished rate of decay of these cells, as in the other two models. Interestingly, none of the three models shows a strong sensitivity of virus to the concentration of IFN in the system.

All models must make some simplifying assumptions, and thus, no models are fully accurate in their representation of the host response to pre-stimulating IFN-regulating pathways. While these models had been analyzed in previous reviews [54], previous work had only shown how these models respond to knockouts of various immune components. Here, we perform a complementary study to test early stimulation as well as increased initial levels of IFN to determine if altering IFN levels can improve patient outcomes. Of the three models studied, only one showed significant impact after early IFN induction. The Saenz model predicts a lower viral peak with increased initial interferon levels and a monotonic decay of the virus over time if interferon is stimulated early. Early available interferon creates a large pool of resistant cells in a short period of time and, because the Saenz model does not allow for loss of resistance in cells over time, resistant cells remain resistant for the rest of the simulation. Thus, the system cannot replenish the source of target cells for the virus to infect and the virus concentration decreases steadily. However, this is not a realistic way to represent immune dynamics, as epithelial cells certainly lose their viral resistance over time.

The other two models do not demonstrate this effect of IFN on viral clearance. The Pawelek model is structured such that early administration of interferon-inducing compounds worsens the impact of the virus by creating a secondary rebound of virus in later stages of infection, essentially leading to chronic infection (which is unlikely to be realistic). The Hancioglu model shows no sensitivity to the initial interferon concentration or to the relative timing of infection. Changing the time of the virus infection simply shifts the curves in time but does not change their shape. This model implies that interferon has minimal impact on the host, which does not agree with decades of experimental evidence [29,55,56].

By stimulating an early IFN response in the model, we simulate a host receiving a preventative treatment for IAV infection (e.g. a TLR agonist [22,28]). Dobrovolny *et al*. [54] previously investigated how these models react to IFN suppression post-infection, which may suffice to simulate a steroid treatment for influenza as steroids are known to downregulate IFN signaling [20,57]. The Hancioglu and Pawelek models predict that IFN has a significant impact on viral clearance when completely removed from the system because no resistant cells are created and the population of susceptible cells remains high for a longer period of time [54]. In the Saenz model, however, removing IFN does not yield this effect, as cells in this model cannot lose resistance. Therefore, these models may not accurately represent the effect of IFN pre-stimulation for influenza, as they make many simplifying assumptions about the role of IFN in the host immune response to IAV infection.

Currently, we do not have sufficient experimental or computational evidence to support a recommendation for IFN pre-stimulation or corticosteroid treatment post-infection. Few references exist showing steroid treatment of IAV-infected humans [9,58–60], and those few have not shown significant impacts on mortality rate [61]. For many years, physicians turned toward high doses of steroids, though recent research suggests that lower doses are more effective [61,62]. It is quite possible that steroid treatment could be effective in humans, but the timing and magnitude of the drug has not yet been optimized. Tan *et al*. [47] have shown Pam2Cys, a TLR-2 agonist, can instigate an inflammatory response even in the absence of an antigen. Mice pre-treated with Pam2Cys were protected from H1N1 virus for up to 7 days post-treatment. Pre-stimulation of TLR-3 by polyinosinic:polycytidylic acid (poly IC) has also shown promise in protecting mice against H5N1 and H3N2 [48]. TLR-3 pre-stimulation has been effective in protecting rhesus monkeys from yellow fever [63]. While IFN-prestimulating compounds have been very promising in animal models, they are still in the early phases of drug development [46]. Nonetheless, the effects of IFN pre-stimulation has been well established and the dynamics induced by pre-stimulation are highly valuable for mathematical model discrimination.

While the models presented do capture many aspects of the immune response to IAV infection, more experimental data is needed to improve the characterization of IFN-regulated immune dynamics. Shinya *et al*. [64] demonstrated the IFN pre-stimulation from 12h to 3 days pre-infection improved survival to IAV-infected mice, but a more thorough dosing range and high temporal resolution of the data are needed to improve model development and validation.

This review has shown that simply creating a population of virus-resistant cells is not sufficient to model the impact of IFN on control of virus replication. This is the mechanism by which many current published models, including the three covered in this paper, incorporate the effect of IFN on the immune response. For a truly accurate mathematical model, the model structure should be able to simulate known qualitative behaviors as well as reproduce the quantitative data used to tune the model parameters. The models used in this review do not include a mechanism by which IFN levels can be sustained if the virus is not present in the system (see Figures 5b, 6b, and 7b). The pre-stimulation is thus ineffective because IFN decays monotonically until the virus is administered days later. Additional cellular sources of IFN production, such as monocytes, may be necessary for a biologically accurate ODE model. The Hancioglu model does include a term for macrophage-derived IFN production, but macrophages are only induced to produce IFN if the virus has been introduced to the system, so this model cannot sustain increased IFN concentration in the absence of pathogen. The Pawelek and Saenz models only contain infected epithelial cell production of IFN.

Future ODE models of influenza infection should include a better representation of innate immunity, and possibly more interactions of IFN with other components in the model, to accurately portray the impact of IFN on the system as a whole. Rather than reliance on the creation of virus-resistant cells to simulate the effect of IFN on the host, IFN could be used to directly diminish the replication rate of the virus, similar to a model proposed by Baccam et al. [40]. Alternatively, IFN could be used to lower the infectivity of the virus and slow the creation of infected epithelial cells. These models could then be used to test the protection conferred by IFN pre-stimulation seen in many murine models of influenza A virus infection [21,22,28,33,65,66].

## Acknowledgments

The authors acknowledge funding support from the University of Pittsburgh’s Swanson School of Engineering and the University of Pittsburgh’s Central Research Development Fund (CRDF).

## Author Contributions

E.M. and J.E.S. conceived and designed the experiments; E.M. performed the experiments; E.M., E.A., and J.E.S. analyzed the data; E.M., E.A., and J.E.S. wrote the paper.

## Appendix A

**Table A1.** Parameter definitions and values for the Pawelek model.

| Parameter | Definition | Value | Units |
| --- | --- | --- | --- |
| $\beta$ | Infectivity of virus | $4.7 \times 10^{-5}$ | (RNA copy)<sup>-1</sup><br>ml NS day<sup>-1</sup> |
| $\phi$ | Rate of production of virus-resistant epithelial cells | 0.33 | (IFN fold<br>change)<sup>-1</sup> day<sup>-1</sup> |
| $\rho$ | Rate of loss of resistance in epithelial cells | 2.6 | day<sup>-1</sup> |
| $\delta$ | Death of infected epithelial cells | 2 | day<sup>-1</sup> |
| $\kappa$ | Rate of infected cell clearance by natural killer cells | 4.2 | (IFN fold<br>change)<sup>-1</sup> day<sup>-1</sup> |
| $p$ | Replication rate of virus | $5.3 \times 10^{-3}$ | (RNA copy)<sup>-1</sup><br>day<sup>-1</sup> cell<sup>-1</sup> |
| $c$ | Clearance of virus by immune system | 16 | day<sup>-1</sup> |
| $q$ | Rate of production of interferon by infected cells | $9.6 \times 10^{-10}$ | (IFN fold<br>change) day<sup>-1</sup><br>cell<sup>-1</sup> |
| $d$ | Rate of depletion of interferon | 1.99 | day<sup>-1</sup> |

**Table A2.** Parameter definitions and values for the Saenz model.

| Parameter | Definition | Value | Units |
| --- | --- | --- | --- |
| $\beta$ | Infectivity of virus | $1.4 \times 10^{-4}$ | (RNA copy)<sup>-1</sup><br>ml NS day<sup>-1</sup> |
| $\phi$ | Rate of production of virus-resistant epithelial cells | 56 | (IFN fold<br>change)<sup>-1</sup> day<sup>-1</sup> |
| $k_1$ | Rate of production of unprotected infected cells | 2 | day<sup>-1</sup> |
| $\delta$ | Death of infected epithelial cells | 2 | day<sup>-1</sup> |
| $m$ | IFN-reduced infectivity | 1 | unitless |
| $k_2$ | Eclipse phase of interferon-protected infected cells | 2 | day<sup>-1</sup> |
| $\alpha$ | Rate at which partially-resistant cells become fully resistant | 4 | day<sup>-1</sup> |
| $p$ | Replication rate of virus | $1.4 \times 10^{-5}$ | (RNA copy)<sup>-1</sup><br>day<sup>-1</sup> cell<sup>-1</sup> |
| $c$ | Clearance of virus by immune system | 5.2 | day<sup>-1</sup> |
| $q$ | Rate of production of interferon by infected epithelial cells | $5 \times 10^{-10}$ | (IFN fold<br>change) day<sup>-1</sup><br>cell<sup>-1</sup> |
| $d$ | Rate of depletion of interferon | 6.8 | day<sup>-1</sup> |
| $n$ | IFN-reduced production | 1 | unitless |

**Table A3.** Parameter definitions and values for the Hancioglu model.

| Parameter | Definition | Value |
| --- | --- | --- |
| $\gamma_{HV}$ | Rate of epithelial cells infected by virus | 0.34 |
| $a_R$ | Loss of resistance to infection | 1 |
| $a_I$ | Death of infected cells | 1.5 |
| $b_{HD}$ | Rate of regeneration of epithelial cells | 4 |
| $\alpha_V$ | Nonspecific clearance of virus | 1.7 |
| $b_{IE}$ | Clearance of infected cells by effector cells | 0.066 |
| $b_{FH}$ | Rate of binding of IFN to epithelial cells | 17 |
| $\gamma_V$ | Rate of virus secretion by infected cells | 510 |
| $\gamma_{VA}$ | Rate of antibody neutralization of virus | 619.2 |
| $\gamma_{VH}$ | Rate of adsorption of virus by infected cells | 1.02 |
| $a_{V1}$ | Nonspecific clearance of virus | 100 |
| $a_{V2}$ | Nonspecific clearance of virus | 23000 |
| $b_{MD}$ | Stimulation of antigen presenting cells by dead cells | 1 |
| $b_{MV}$ | Stimulation of antigen presenting cells by virus | 0.0037 |
| $a_M$ | Death of antigen presenting cells | 1 |
| $b_F$ | IFN production by antigen presenting cells | 250000 |
| $c_F$ | IFN production by infected cells | 2000 |
| $b_{HF}$ | Rate of production of virus-resistant cells | 0.01 |
| $a_F$ | Natural decay of IFN | 8 |
| $b_{EM}$ | Production of effector cells | 8.3 |
| $b_{EI}$ | Death of effector cells through interaction with infected cells | 2.72 |
| $a_E$ | Natural death of effector cells | 0.4 |
| $b_{PM}$ | Production of plasma cells | 11.5 |
| $a_P$ | Death of plasma cells | 0.4 |
| $b_A$ | Production of antibodies by plasma cells | 0.043 |
| $\gamma_{AV}$ | Rate at which antibodies bind to virus | 146.2 |
| $a_A$ | Natural decay of antibodies | 0.043 |
| $r$ | Rate of compatibility of antibodies and virus | $3 \times 10^{-5}$ |

